# Circadian clock features define novel breast cancer subtypes and shape drug sensitivity

**DOI:** 10.1101/2024.10.07.617001

**Authors:** Carolin Ector, Jeff Didier, Sébastien De Landtsheer, Malthe S. Nordentoft, Christoph Schmal, Ulrich Keilholz, Hanspeter Herzel, Achim Kramer, Thomas Sauter, Adrián E. Granada

## Abstract

The circadian clock regulates key physiological processes, including cellular responses to DNA damage. Circadian-based therapeutic strategies optimize treatment timing to enhance drug efficacy and minimize side effects, offering potential for precision cancer treatment. However, applying these strategies in cancer remains limited due to limited understanding of the clock’s function across cancer types and incomplete insights into how the circadian clock affects drug responses. To address this, we conducted deep circadian phenotyping across a panel of breast cancer cell lines using two complementary reporters. Observing diverse circadian dynamics, we developed metrics to assess circadian rhythm strength and stability. This led to the identification of four distinct circadian-based phenotypes in breast cancer: functional, weak, unstable, and dysfunctional clocks. Furthermore, we demonstrate that the circadian clock plays a critical role in shaping pharmacological responses to various anti-cancer drugs and identify circadian features that accurately predict drug sensitivity. Collectively, our findings establish a foundation for advancing the use of chronotherapeutic strategies in breast cancer treatment, expanding their potential application to improve therapeutic outcomes in breast cancer.

## Introduction

In alignment with the solar 24-hour day-night cycle, the circadian clock regulates essential physiological and behavioral processes in almost all organisms. In mammals, a hierarchical structure coordinates rhythmic activities across both whole-organism and cellular levels (Chaix *et al*, 2016; Golombek *et al*, 2014), with the master clock residing in the suprachiasmatic nucleus (SCN) and individual peripheral clocks oscillating across tissue types (Chaix *et al*., 2016). Remarkably, about 40% of all genes are rhythmically expressed in a tissue-specific manner (Zhang *et al*, 2014), influencing various biological functions such as metabolism (Neufeld-Cohen *et al*, 2016), cell growth (Chakrabarti & Michor, 2020), immune responses (Scheiermann *et al*, 2013), and DNA repair (Sancar *et al*, 2010), thereby maintaining cellular balance.

At its core, the circadian mechanism is composed of transcriptional-translational feedback loops (TTFLs) (Takahashi, 2017). These TFFLs consist of the CLOCK and BMAL1 transcription factors, which induce the transcription of *Per1/2/3* and *Cry1/2* genes through the binding to the respective promoter regions. In turn, PER and CRY proteins inhibit the binding of CLOCK/BMAL1, and consequently their own transcription. In a complementary second feedback loop the transcription of *Bmal1* is rhythmically regulated by REV-ERBα/β repressor and RORα/β/γ activator proteins, whose transcription is likewise dependent on the CLOCK/BMAL1 transactivation complex (Chaix *et al*., 2016; Takahashi, 2017).

Environmental factors are crucial in maintaining the synchronization of circadian rhythms in our body. Misalignments of this synchronization, for example through prolonged exposure to shift work, is associated with various diseases, including cancer (Sulli *et al*, 2019) – the second-leading cause of death worldwide (https://www.who.int/health-topics/cancer, 2022). In-depth studies have further shown how the disturbance of the circadian clock, through either genetic mutations or the suppression of core clock genes is directly linked to the progression of cancer and poorer survival rates (Ye *et al*, 2018). The clock’s influence extends to cancer therapy, affecting the efficacy and toxicity of treatments in a time-of-day-dependent manner (Lee *et al*, 2021; Ye *et al*., 2018). Despite its high medical importance, the distinct role of the circadian clock in cancer remains poorly understood, highlighting the need for further research to optimize treatment strategies by circadian rhythms.

To illuminate the circadian clock’s role in cancer, our study focuses on breast cancer, the most prevalent cancer among women and the second most common overall (Bray, 2024). Breast cancer is categorized into four main subtypes, distinguished by the cell phenotype (basal or luminal) and specific biomarkers, including progesterone and estrogen receptors, as well as HER2-receptor overexpression (Sotiriou *et al*, 2003) (**Fig. 1A**). Within the highly heterogeneous and aggressive triple-negative breast cancer (TNBC) subtype lacking all three biomarkers, Lehmann et al. identified four molecular TNBC subtypes: basal-like 1 (BL1) and 2 (BL2), mesenchymal (M) and immunomodulatory (IM) (Lehmann *et al*, 2011). While the presence of circadian rhythms have been shown for selected breast cancer models (Lellupitiyage Don *et al*, 2019; Li *et al*, 2024), the general assumption is that more aggressive breast cancer subtypes including TNBC exhibit disrupted circadian rhythms (Lellupitiyage Don *et al*., 2019; Li *et al*., 2024; Rida *et al*, 2019). In this study, we aim to define novel circadian clock-based phenotypes within breast cancer and to uncover how circadian properties relate to drug sensitivity and tumor growth (**Fig. 1B**). For this, we employ a comprehensive set of data analysis methods to deep-phenotype the circadian clock (**Fig. 1C–E**), and to collect information on cellular growth dynamics (**Fig. 1F**), drug responses (**Fig. 1G**), and genomic properties of circadian clock genes (**Fig. 1H**). Subsequently, we integrate our acquired cellular parameters to identify cell clusters and establish novel circadian-based subtypes within breast cancer (**Fig. 1I**). Finally, we investigate how drug responses depend on the circadian clock, identifying several drugs that could be further explored for their potential in chronotherapeutic approaches. Altogether, our approach sets the stage for a deeper understanding of distinct tumor-specific circadian clocks and how they shape cancer therapy outcomes.

**Figure 1.**
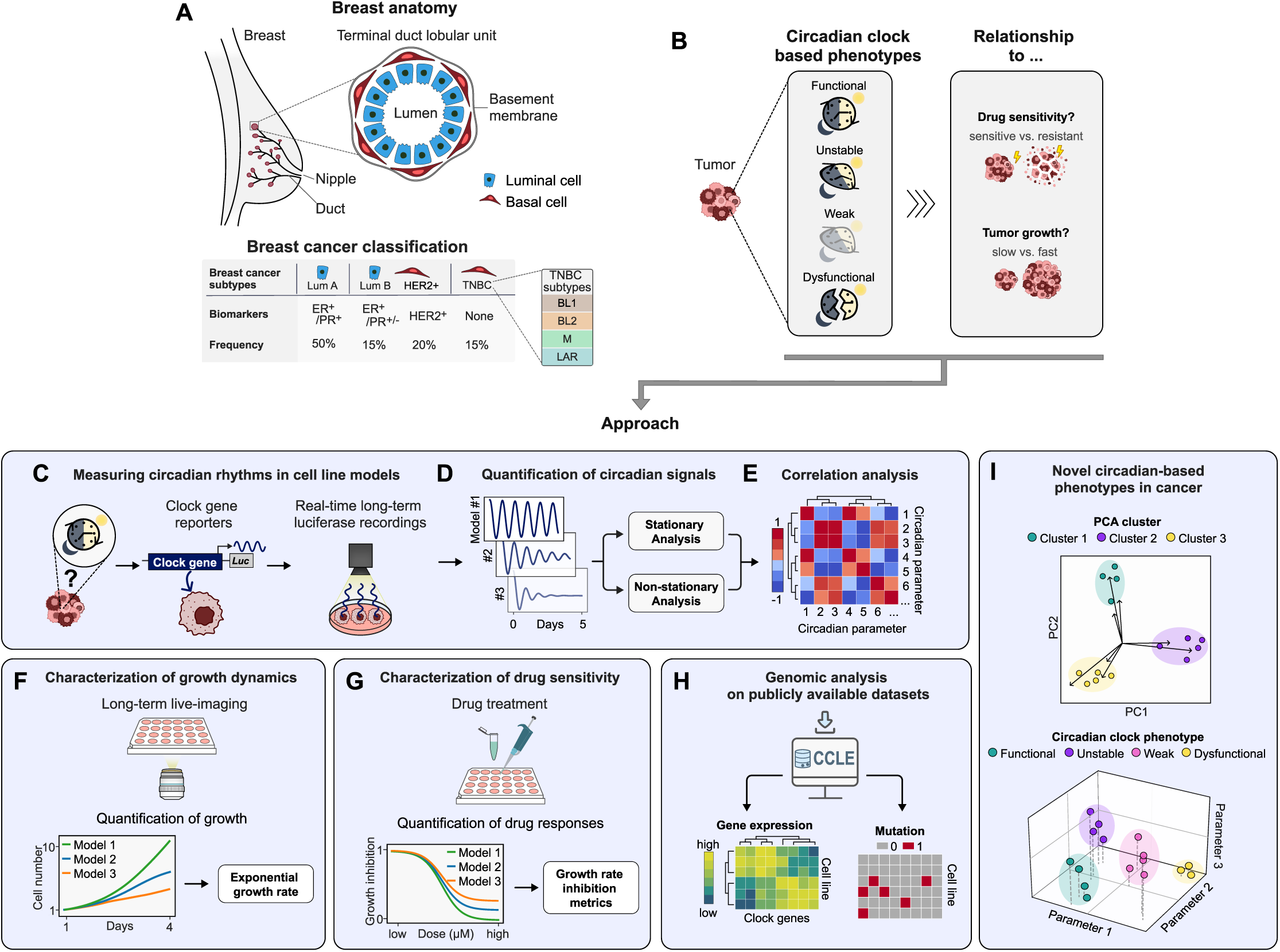
Definition of circadian-based breast cancer subtypes by deep phenotyping. **(A)** Classification of breast cancer based on cancer cell phenotype and biomarker expression status. Approximate frequency among all breast cancer cases indicated in percentage. -, negative; +, positive; ER, estrogen receptor; PR, progesterone receptor; HER, human epidermal growth factor receptor 2; LumA/B, Luminal A/B; TNBC, triple-negative breast cancer, BL1/2, basal-like 1/2; M, mesenchymal; LAR, luminal androgen receptor. (B) Aim of our study to identify circadian-based phenotypes in breast cancer and investigate the role of circadian clock features for tumor growth and drug sensitivity. **(C)** Cellular circadian rhythms were measured by long-term luciferase (Luc) recordings of clock gene reporter cell lines. **(D)** Circadian signals were quantified by stationary and non-stationary analysis approaches and **(E)** related to each other by correlation analysis to define a set of representative circadian metrics. (F) Long-term live imaging was employed to capture growth dynamics and **(G)** drug sensitivities across various cell line models, followed by the parametrization of representative metrics. **(H)** Characterization of genomic circadian profiles of the cell models tested, utilizing publicly available datasets. (I) Definition of circadian-based phenotypes through the integration of multiple cellular parameters.

## Results

### Deep circadian phenotyping reveals variability in clock strength across breast cancer models

The role of the circadian clock in cancer progression and treatment is gaining increasing interest, yet the degree of circadian rhythmicity and intrinsic timing profiles for various cancer subtypes remains largely unclear. To address this, we utilized high-resolution circadian clock recordings alongside a range of time-series analysis techniques to effectively measure and characterize circadian rhythms in cancer models. Building on our previously described deep-circadian phenotyping approach (Ector *et al*, 2024), which is comprehensively detailed in **Box 1**, we expanded our dataset by incorporating additional cell line models and novel analysis methods.

To obtain an initial estimate of the periodic quality of the time-series, we commenced our deep circadian phenotyping approach with autocorrelation (AC) analysis. Screening of 19 different cancer and non-cancer cell lines with the majority classifying as the highly heterogeneous and aggressive TNBC, we identified a broad spectrum of rhythmicity indices and period lengths, mostly centered around the 24-hour human circadian cycle. Interestingly, rhythmicity indices below 0.3, indicative for weaker rhythms, often deviated from 24 hours, whereas higher indices aligned with the circadian cycle (**Fig. 2A**). These results indicate a considerable variability of circadian phenotypes within the cell lines tested, highlighting the need to further dissect the circadian characteristics.

**Figure 2.**
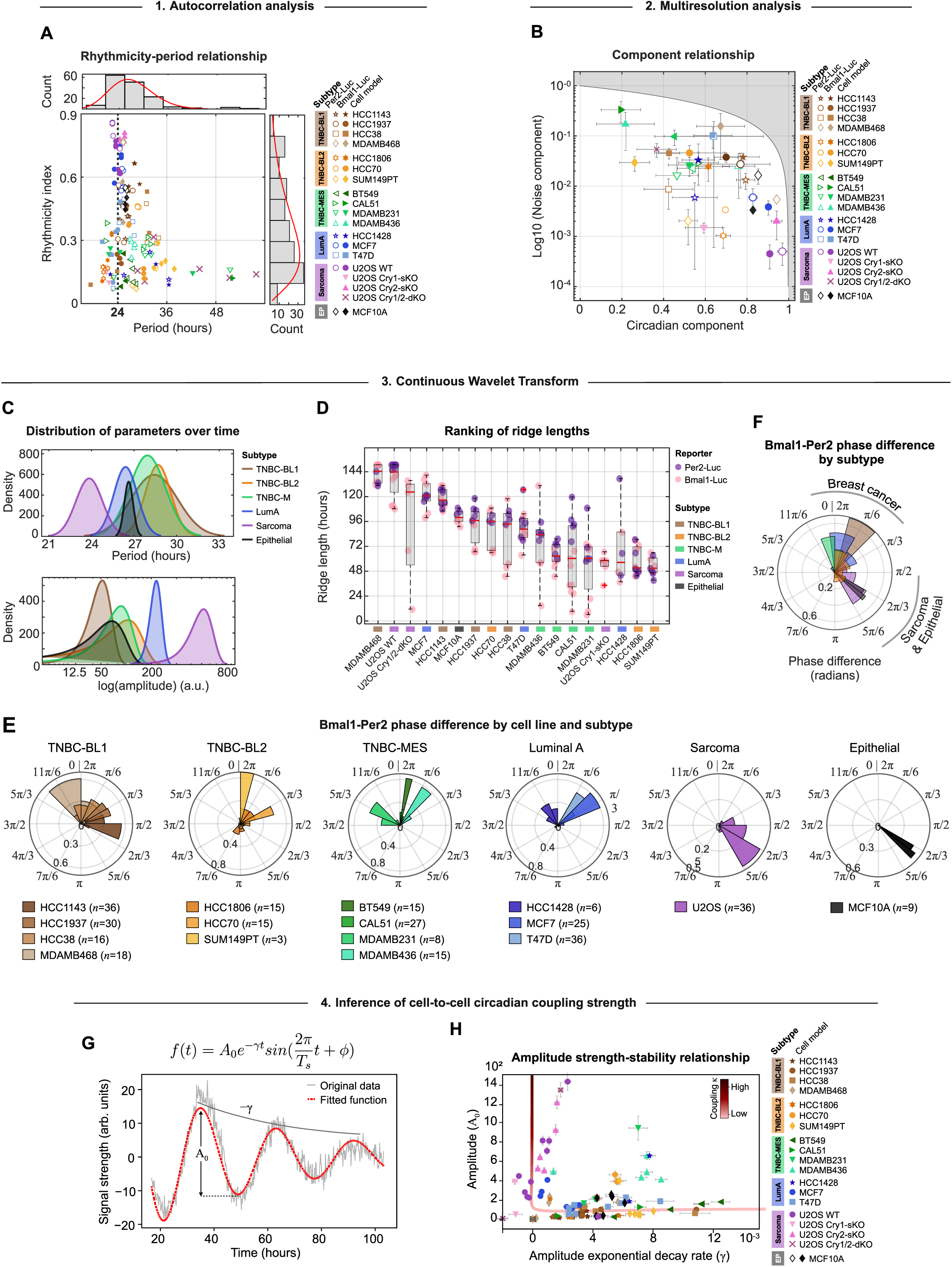
Deep phenotyping of circadian rhythms in breast cancer cell line models. **(A)** Relationship between rhythmicity indices and periods for various cell line models, distinguished by markers and colored by subtype (n=141 samples represented as individual measurements). Dashed line at period=24 hours. Gamma distribution of periods and rhythmicity indices shown in histograms in the top and left panel, respectively. **(B)** Relationship between normalized noise and circadian components across cell line models, determined by multiresolution analysis. Cell models are color-coded by subtype and distinguished by markers. Shaded area covers unattainable range. Data represents the mean±s.d. of 6 samples per reporter (n=2 biological replicates a technical triplicates or duplicates [HCC1806 Per2-Luc, n=5]). *n=3* for MCF10A and MDAMB468-Per2-Luc (single experiment). **(C)** Normal distribution of subtype-averaged and time-resolved periods (top panel) and amplitudes (bottom panel) (n=1-4 cell lines models per subtype). Before calculating subtype averages, all available replicates per cell model were averaged (n=3-6, see **B** for exact numbers) and used as input. **(D)** Boxplot of CWT ridge lengths from *Bma/1-* and Per2-signals of specified cell models, with the bottom and top edges of the boxes representing the 25^th^ and 75^th^ percentiles, respectively. Extending whiskers represent data points within 1.5 times the interquartile range from lower and upper quartile. Median values are marked by red horizontal lines, and outliers by red crosses. n=4-12 samples per cell line (see **B** for exact numbers). **(E)** Subtype-specific polar histograms of *Bma/1-Per2* phase differences over time, averaged and color-coded by cell line. The number *n* of Bma/1-Per2-combinations per cell line is indicated next to the cell line name. Polar histograms were normalized by probability. **(F)** Polar histogram of *Bma/1-Per2* phase differences over time, averaged and colored by subtype (n=1-4 cell line models). In the polar histograms in **E** and **F** 2n corresponds to one full circadian cycle. **(G)** Example of fitting signals to an exponentially decaying sinusoidal model with constant periods to deduce initial amplitudes (Ao) and amplitude decay rates (y). **(H)** Relationship between initial signal amplitudes and amplitude decay rates across cell line models. Cell models are color-coded by subtype and distinguished by markers. Error bars indicate fitting errors. The L-shaped line represents the simulated coupling strength (K).

To quantify the proportion of distinct frequency components, present in each signal, we employed multiresolution analysis (MRA). This process segmented the detrended signal into four frequency bands: noise (1–4 hours), ultradian (4–16 hours), circadian (16–32 hours), and infradian (32–256 hours) (see **Methods**). Relating the circadian to the noise component revealed a spectrum of signal-to-noise ratios across cell lines, where higher circadian signals corresponded to lower noise, indicating circadian signal robustness, and vice versa. Alongside the osteosarcoma cell line U-2 OS, which is well-studied for its functional circadian clock (Baggs *et al*, 2009), high signal-to-noise ratios were found for the epithelial MCF10A, the LumA-MCF7 and the TNBC-MDAMB468 cell line (**Fig. 2B**). Interestingly, a knockout of *Cry2* in the U-2 OS cell line alone did not essentially alter the signal-to-noise ratio. In contrast, the knockout of *Cry1* or both, *Cry1* and *Cry2*, decreased the circadian component substantially by 34.9% (*p=*5.3×10^−6^) and 59.8% (*p=*8.9×10^−8^), while significantly enhancing the noise component by 3.3-fold (*p=*9.0×10^−5^) and 120.1-fold (*p=*1.2×10^−5^), respectively.

### Clock dynamics vary within subtypes of breast cancer

After evaluating the strength and composition of rhythmic structures within our signals, we analyzed their dynamic nature through continuous wavelet transform (CWT). This approach visualizes the signal in a CWT power spectrum heatmap, showcasing the range and relative power of signal components over time within a specified period range. By tracking the main oscillatory component (“*ridge*”), CWT effectively reveals non-stationary and time-dependent features, such as dynamic periods and amplitudes (see **Fig. S1A** and **Methods**).

Aggregating the distribution of weighted mean periods and amplitudes per subtype, we noted variability of these parameters over time, particularly for the three TNBC subtypes (**Fig. 2C**; see **Fig. S1B** and **Methods** for weighting approach). Furthermore, we revealed varying prevailing periods and amplitudes for each subtype where only the osteosarcoma subtype showed a distinct period peak at approximately 24 hours, combined with the highest amplitudes, while the prevailing periods for the other subtypes were mainly prolonged (**Fig. 2C**). Focusing on the breast cancer subtypes, a clear differentiation in circadian properties was visible, where the more aggressive TNBC subtypes exhibited longer periods and lower amplitudes than the LumA subtype (**Fig. 2C**). Though, when considering median CWT periods and amplitudes of each cell model, we found considerable within-subtype diversity, indicating that individual cancer tissue types exhibit a spectrum of circadian clock phenotypes (**Fig. S1C**). This within-subtype diversity extends to the ridge lengths, a measure of circadian clock strength, where longer and continuous ridges denote robust signals and shorter, discontinuous ridges indicate weaker ones (**Fig. 2D**). Here, only TNBC-M cell models ranked alongside of each other, whereas cell models of the LumA and basal-like TNBC subtypes displayed considerable variability in ridge lengths.

Building on the identified temporal circadian dynamics within and across cancer subtypes, we next assessed *Bmal1*-*Per2* phase differences over time. Consistent with the previously discussed circadian parameters, we observed a spectrum of phase differences across and within tissue types (**Fig. 2E** and **2F**). Despite this variability, a notable pattern emerged in which non-cancerous and osteosarcoma tissues consistently showed phase differences around 2π/3, an eight-hour lag in a 24-hour cycle, in contrast to breast cancer subtypes, which tended to have no phase difference (2π) (**Fig. 2F**).

Circadian clocks in breast cancer cell lines may exhibit either self-sustained or damped oscillatory behaviors, a distinction that depends on the degree of intercellular coupling. Aiming to infer intercellular circadian coupling strength, we next analyzed signal amplitudes and corresponding exponential decay rates. Inspired by previous studies (Del Olmo *et al*, 2024; Finger *et al*, 2021; Guenthner *et al*, 2014), we used a network of identical Poincaré oscillators to model individual cells within a tissue, featuring a constant coupling strength (κ) and periods averaging a 24-hour cycle. By varying the coupling strength, we observed distinct signal patterns, with strong coupling leading to amplified and self-sustained oscillations, while weak coupling results in a damped oscillation pattern (**Fig. S2**).

We then fitted our experimental data to an exponentially decaying sinusoidal function (**Fig. 2G** and **Methods**) to determine initial amplitudes (A_0_) and amplitude decay rates (*γ*). A comparative analysis unveiled an L-shaped trend in both, the simulated and experimental datasets, facilitating the approximation of coupling strengths in experimental data (**Fig. 2H**).

Here, stronger intercellular coupling was identified for signals with high amplitudes and slow decay rates and vice versa. These findings align with previous studies, which demonstrated that a reduction in coupling strength results in increased variance, disrupting both the summed amplitude and period (Del Olmo *et al*., 2024; Guenthner *et al*., 2014). Furthermore, focusing on U-2 OS cells, we observed a significant decrease in coupling strength in the circadian knockout cell lines compared to the wild-type (**Fig. 2H**), which is consistent with predictions from earlier research (Del Olmo *et al*., 2024).

These insights into the dynamics of circadian clock features, together with the broad range of circadian rhythm strengths previously described, underscore the heterogeneity of circadian regulation within cancer tissues. Such within subtype-diversity motivated us to establish a novel circadian-based classification that could potentially inform chronotherapeutic strategies.

### Establishing a novel circadian-based subtyping in breast cancer

After extracting various circadian clock parameters from the different cell lines, we next investigated whether circadian clock-based clusters are present among these lines and how this clustering relates to their classical subtypes. Considering the established molecular connections between the circadian clock and cell cycle (Feillet *et al*, 2015; Gonze, 2024; Gutu *et al*, 2024), we integrated cellular growth rates into our analysis, which we determined by exponential curve fitting of long-term growth data (**Fig. S3**). To streamline our extensive dataset, we initially assessed the relationships between the parameters by calculating pairwise Pearson correlation coefficients, aggregating data from all replicates per cell line and from both circadian reporters. In line with our expectations, we discovered several strong correlations as well as redundancies across the dataset (**Fig. S4A**), allowing us to narrow down the circadian dataset to key parameters shown in **Fig. 3A**. We observed significant positive correlations between the different clock strength metrics, highlighting consistency across the different signal analysis approaches. In contrast, clock-strength parameters were mostly negatively, yet weaker, correlated with the growth rate, demonstrating accelerated growth in cell models with weaker circadian rhythms, whereas parameters representative of clock instability such as period and phase difference variations over time were positively correlated with growth (**Fig. 3A** and **Fig. S4A**).

**Figure 3.**
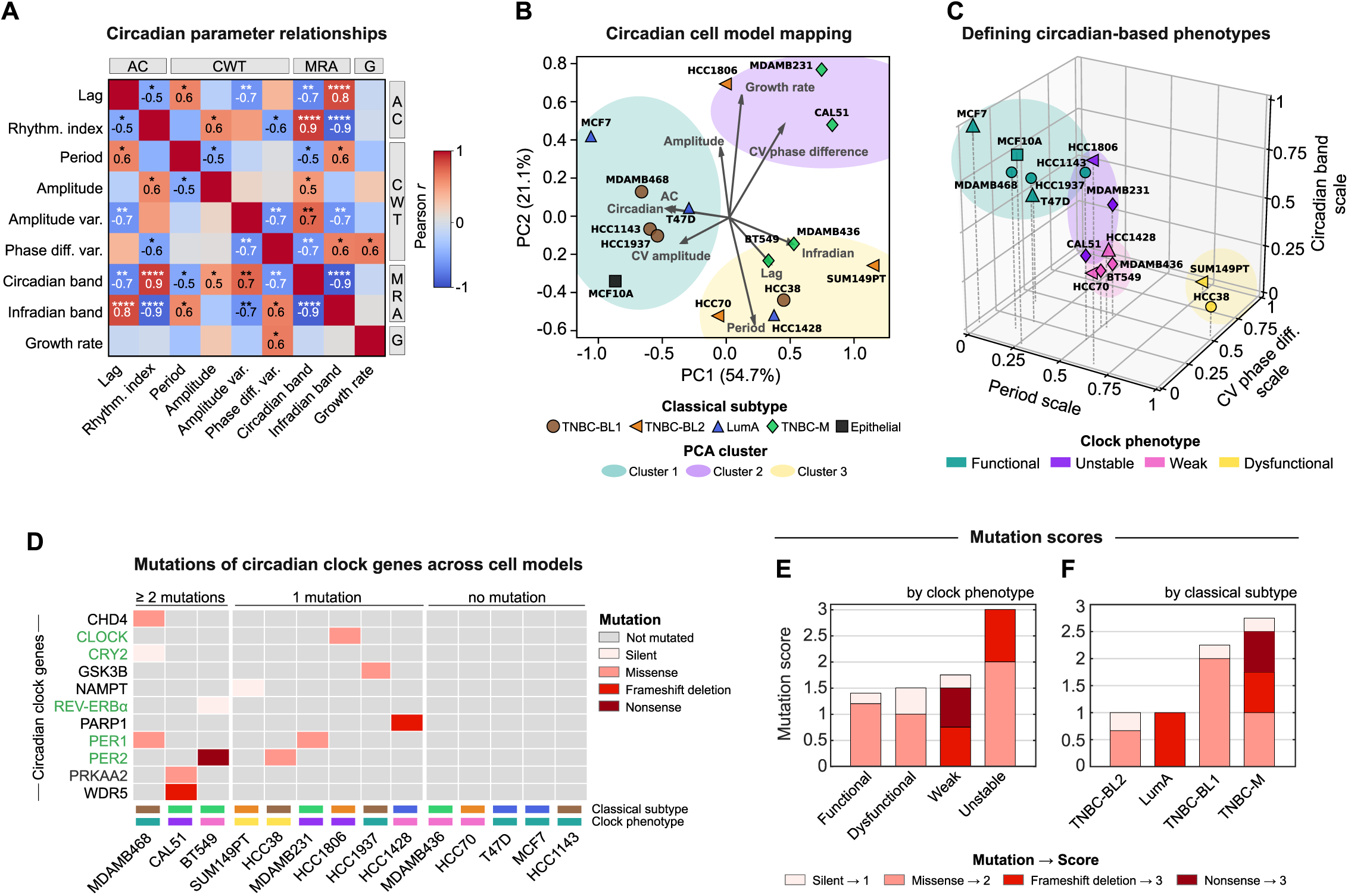
Circadian clock features define circadian-based subtypes in breast cancer. **(A)** Pearson correlation coefficients between selected *Bmal1-* Per2-averaged circadian features and growth rates across all breast cancer cell line models and the epithelial MCF10A cell line (n=15 cell lines). Parameters are categorized by their approach of calculation: AC, autocorrelation; CWT, continuous wavelet transform; MRA, multiresolution analysis; G=growth. Displayed are statistically significant correlation values, where *, **, ***, and **** indicate p-values < 0.05, 0.01, 0.001, and 0.0001, respectively. (B) Principal component analysis (PCA) biplot of the indicated circadian and growth parameters, displaying the distribution of 15 cell line models across the first two principal component scores. Cell models are depicted by different markers indicating their classical subtype. Color-coded clusters are outlined around closely positioned cell models, based on visual inspection. The variance (PC loadings) explained by each component is expressed in percentage. CV=coefficient of variation. **(C)** Three-dimensional space showing the k-means clustering *(k=4)* of the indicated cell line models. Discretization was based on three representative, min-max-scaled parameters from each PCA-cluster defined in subplot B. Novel clock phenotype clusters are color-coded. Classical subtypes are denoted with different markers (refer to subplot B for key). Within cluster sum of squares=0.79, cluster silhouette score=0.45. CV, coefficient of variation. (D) Hierarchical clustering of cell lines by mutational burdens in core circadian clock (green) and circadian clock associated genes (black). The heatmap illustrates the mutation type by cell line and gene. Color-coded rectangles above the x-labels indicate the classical subtype and clock phenotype (refer to subplot **B** and **C** for color-coding). See Supplementary Table 1 for full list of investigated circadian genes. **(E)** Ranking of averaged mutation scores by clock phenotype or **(F)** by classical subtype, with stacked bars showing the mutation type proportions contributing to the final score, n=2-4 cell models; for exact numbers of cell line models per group refer to panel B.

Upon refinement of our dataset, we employed principal component analysis (PCA) to investigate the presence of circadian-based clusters within the breast cancer and epithelial cell line panel. PCA showed the formation of three distinct circadian-based clusters where the respective cell models share characteristics of circadian clock or growth dynamics (**Fig. 3B**). Notably, most of the variability in the dataset was captured by the first two principal components alone, which accounted for 54.3% and 21.8% of the variance, respectively. Ranking of parameters by their corresponding PC loading, revealed that the highest influence on the observed clustering for the first principal component comes from the infradian and circadian components, underlining their potential as key markers in the circadian characterization of the cell models (**Fig. S4B**). Interestingly, the circadian-based grouping deviated from the classical subtyping, providing a novel perspective on the subtyping of these cell models.

Since the identified PCA clusters do not directly translate into circadian phenotypes, we selected one key circadian parameter of each of the identified PCA clusters and performed K-means cluster analysis (**Fig. 3C**). In detail, we focused on the oscillation period, the *Bmal1-Per2* phase difference variability over time for clock stability, and the circadian component as a measure of clock strength. As a result, we successfully classified the cell models into four circadian phenotypes based on their circadian clock functionality: functional, unstable, weak, or dysfunctional (**Fig. 3C**). The functional phenotype, marked by a high circadian component and stable phase difference, suggests a strong and consistent circadian rhythm, and is composed of the epithelial MCF10A, two LumA, and two TNBC-BL1 cell line models. In contrast, the dysfunctional phenotype, comprising of two cell models only, is characterized by instable phase differences over time, long periods and low circadian bands. The intermediate phenotypes, weak and unstable, display decreased rhythm strength and stability, respectively (**Fig. 3C**). Consistent with the clustering, cell models sharing similar circadian clock phenotypes were mapped in proximity using Uniform Manifold Approximation and Projection (UMAP), which enabled the visualization of all key circadian parameters in a reduced dimensional space (**Fig. S4C**).

In summary, we introduced a new phenotypic framework that categorizes breast cancer cell lines according to their circadian phenotypes. This approach provides a complement to traditional subtyping methods, potentially enhancing the evaluation of cancer models for circadian-based therapeutic strategies.

### Genetic profiles of circadian-based phenotypes

To evaluate the role of circadian genetic profiles in the newly identified circadian-based phenotypes, we extracted the mutation and gene expression profiles of 16 core clock genes and 44 circadian clock-related genes from the cancer cell line encyclopedia (Barretina *et al*, 2012) (**Supplementary Table 1**). We specified core clock genes as those essential for regulating the circadian TTFLs and for which any dysregulation or mutation can lead to disrupted circadian rhythms (Ko & Takahashi, 2006; Takahashi, 2017).

We noted varying mutation profiles of the circadian genes across the 14 tested breast cancer cell lines, with the majority of cell models exhibiting at least one mutation, whereas five cell lines showed no mutations in their circadian clock network (**Fig. 3D**). We then categorized mutations based on their impact on the protein sequence, assigning scores from 1 for silent mutations to 3 for highly damaging mutations like nonsense and frameshift mutations. Thereby we could compare the clock phenotypes by their circadian mutation burden, revealing that the unstable clock phenotype exhibits the highest mutation score while the functional phenotype had the lowest one (**Fig. 3E**). Notably, damaging mutations appeared exclusively in the unstable and weak clock phenotypes, suggesting a dysregulation of the circadian clock mechanism. Contrary to our expectations, the dysfunctional clock phenotype shared a similar low mutation burden with the functional phenotype, indicating that the number and severity of mutations alone may not fully explain clock functionality, and that additional factors might play a role for defining the underlying clock phenotypes (**Fig. 3E**). When analyzing the mutation scores across classical cancer subtypes, we found that the TNBC-mesenchymal subtype had the highest mutation burden, while TNBC basal-like 2 had the lowest one (**Fig. 3F**). We also identified a tendency for less damaging mutations in basal-like subtypes, whereas Luminal A and TNBC-mesenchymal subtypes exhibited more damaging mutations in the circadian gene panel.

To identify whether expression patterns of core circadian clock genes align with the identified clock phenotypes, we clustered the cell models based on their circadian gene expression dynamics and exposed three overarching clusters (**Fig. S4D**). However, these clusters did not reflect the clock phenotypes but rather showed a distinction between the LumA and the TNBC subtypes. This distinction also emerged when mapping the cell models in a dimensionality-reduced UMAP-space (**Fig. S4E**), indicating that the difference in circadian gene expression dynamics is associated with underlying breast cancer subtypes rather than the intrinsic circadian phenotype.

### Circadian features drive sensitivity profiles to numerous drugs

Due to several interactions between circadian clock genes and therapeutic drug targets, circadian clock-aligned treatments have proved to increase drug effects based on the optimal time-of-day at their administration (Dallmann *et al*, 2014; Lee *et al*., 2021; Ye *et al*., 2018). To unravel the crucial link between circadian features and drug sensitivity, we employed linear regression and linear discriminant analysis on both our own and publicly available drug sensitivity datasets, enabling the screening of various drug classes (**Fig. 4A**; information on drugs listed in **Supplementary Table 2**).

**Figure 4.**
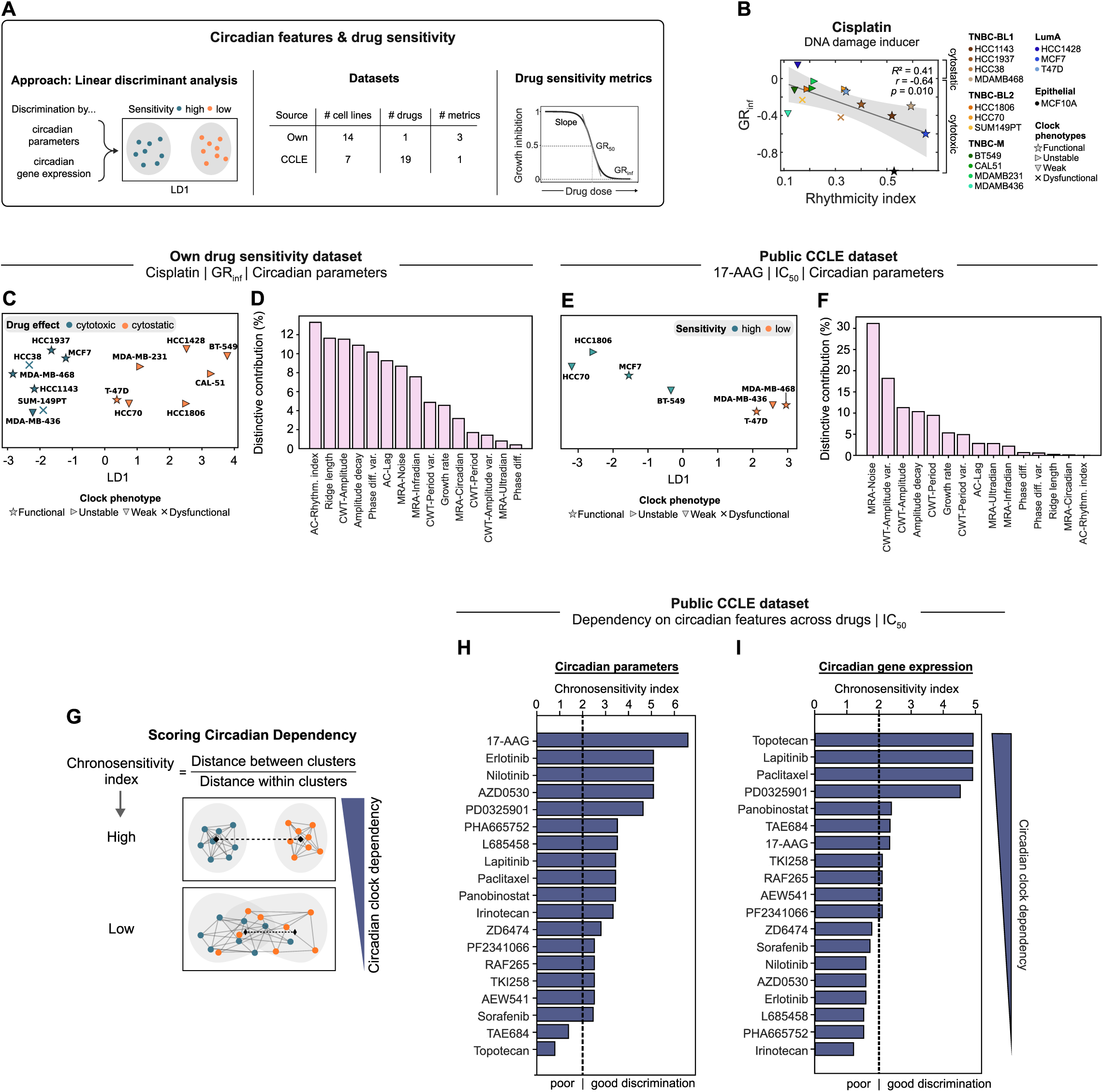
Investigating the role of circadian features in driving drug sensitivity. **(A)** Applied approaches and datasets to identify the role of circadian features in shaping drug sensitivity profiles. **(B)** Linear correlation analysis between cisplatin GR,nf values (absolute drug effects; annoted on the right side of the plot) and the rhythmicity indices assessed by autocorrelation analysis (averaged *Bma/1-Per2* data). Cell models are color-coded and illustrated by their circadian phenotype with distinct markers. Grey-shaded area=95% CI. Model accuracy indicated by R2-values. **(C)** Linear discriminant analysis (LOA) on median-binarized *GR,,1* values to cisplatin, using *Bma/1-Per2* circadian and growth parameters as predictors (n=15 parameters). Cell models exhibiting drug sensitivity values below or above the median are colored in green and orange, respectively. The x-axis represents the first linear discriminant (LD1), which captures the maximum variance between groups. **(D)** Bar plot ranking circadian and growth parameters by their respective contribution to the obtained discriminative information shown in 4d. **(E)** LOA on median-binarized IC50 values to 17-AAG and **(F)** corresponding bar plot ranking the contribution of each parameter to the obtained discriminative information. **(G)** Approach to identify circadian dependencies of drugs (termed as “chronosensitivity index”), calculated from the Between-Cluster-Distance (BCD) to Within-Cluster-Distance (WCD) ratio. The higher the chronosensitivity index, the more accurate the distinction into drug sensitivity groups, hence the higher the dependency on the circadian clock. **(H)** Ranking of chronosensitivity indices of multiple drugs, based on their dependency on circadian oscillation parameters or on (I) circadian gene expression values. The dashed line (y=2) denotes the cut-off for good and poor discrimination within the LOA space.

To account for cell division events during drug perturbation assays and to obtain robust sensitivity metrics, we assessed drug sensitivity using the normalized growth rate inhibition (*GR*) approach (Hafner *et al*, 2016). For this, we expanded the analysis of our drug-response dataset published in Ector et al. (Ector *et al*., 2024). We analyzed a broad spectrum of drugs including DNA synthesis inhibitors (5-FU, doxorubicin), mitosis inhibitors (alisertib, paclitaxel), inhibitors for PI3K (alpelisib), or mTOR (torin2), and DNA damage response (DDR) targeting agents (adavosertib, cisplatin, olaparib). Through pairwise linear correlation analysis, we discovered a significant anti-correlation between the maximal effect of the DDR-inducer cisplatin and the autocorrelation rhythmicity index (*r=*-0.64, *R*^2^=0.41) (**Fig. 4B**). Here, higher rhythmicity was associated with more cytotoxic responses to cisplatin, whereas lower rhythmicity was linked to cytostatic or less toxic responses. This anti-correlation extended to other circadian strength-related parameters such as the amplitude, ridge length and circadian component (**Fig. S5A**). Aligning with these observations, weaker responses to cisplatin were positively correlated with clock instability parameters, such as the variability in periods and phase differences. Besides cisplatin, we identified significant relationships between distinct circadian metrics and *GR*_inf_ values for alpelisib, 5-FU, and alisertib, while *GR*_inf_ values to Olaparib were positively correlated with the growth rate (**Fig. S5A**). Notably, *GR*_inf_ values from alpelisib showed the strongest correlation with circadian parameters (median *r=*0.82), hinting to a role of the circadian clock in driving response phenotypes to the PI3K inhibitor (**Fig. S5B**). Vice versa, we found the highest overall correlation for the circadian amplitude parameter (median *r=*0.74), suggesting a predictive role for maximal drug responses (**Fig. S5C**). Links between circadian parameters and drug response dynamics were further indicated by significant correlations among other drug sensitivity metrics, like the drug concentration at half-maximal effect (*GR*_50_) (**Fig. S5D–F**), and the slope of the dose-response curve (Hill coefficient) (**Fig. S5G–I**).

Despite identifying selected significant correlations between individual clock or growth parameters and drug responses, overall correlations were modest. Given the multilayered nature of the circadian clock, drug sensitivity is likely influenced not by a single circadian factor, but by the collective impact of various circadian features. To investigate this, we utilized linear discriminant analysis (LDA) to discern the cumulative effect of circadian features on categorizing cell models into groups of high or low sensitivity. This method not only assesses the individual contributions of the parameters to the classification but also facilitates the identification of drugs whose effects are likely driven by circadian rhythms (see **Methods**). In our dataset, LDA successfully differentiated cell models based on their cytostatic or cytotoxic responses to cisplatin (**Fig. 4C**). The circadian parameters contributed to varying degrees to this discrimination, with the rhythmicity index and ridge length collectively explaining ∼25% of the separation between the classes, emerging as the most influential discriminative features (**Fig. 4D**). To deepen our understanding of the molecular mechanisms by which the circadian clock modulates cellular responses to drug perturbations, we next investigated the influence of circadian clock gene expression on drug responses. In line with our findings for the circadian parameters obtained from our deep circadian phenotyping approach, the circadian gene set was able to broadly categorize cell models by their responses to cisplatin (**Fig. S6A**, left panel). Examining the individual linear discriminant components of each circadian clock gene revealed a clear ranking of genes in terms of their discriminative importance, with the core clock genes *Rorα* and *Cry2* being the most important contributors, collectively accounting for approximately 34% of the discriminative information (**Fig. S6A**, right panel). The potential of circadian parameters, growth rates and circadian genes to reliably group cell models according to their sensitivity to cisplatin was further identified for *GR*_50_ values and Hill coefficients of the response curves (**Fig. S6B–E**).

To broaden the scope of our analysis, we incorporated publicly available drug sensitivity data from a subset of our circadian-phenotyped cell models. Our analysis revealed that sensitivity to the HSP90 inhibitor 17-AAG was particularly well discerned by circadian parameters (**Fig. 4E**). Among these, factors related to clock stability contributed most to the classification of cell models into high and low sensitivity groups, based on their respective IC_50_ values (**Fig. 4F**). To quantitatively evaluate LDA performances and the discriminative power of circadian features on drug responses, we calculated the ratio of between-cluster-distances (BCD) to within-cluster-distances (WCD), termed as the “chronosensitivity index” of a drug in the following (**Fig. 4G**). Strikingly, the circadian parameters provided a well separation of cell models into sensitivity groups for almost all 19 drugs analyzed (**Fig. 4H**), as indicated by chronosensitivity indices greater than 2, which we defined as a threshold for effective separation (see **Fig. S6F** for a direct comparison of LDA profiles with different chronosensitivity indices). Furthermore, we found that circadian clock gene expression levels could predict the sensitivity patterns of the cell models for a number of drugs (**Fig. 4I**). However, for approximately half of the drugs, the chronosensitivity index fell below the threshold, indicative of poor separation into sensitivity groups. Notably, although circadian parameters did not effectively classify cell models by their sensitivity to the topoisomerase inhibitor topotecan, its chronosensitivity index was substantially higher for the circadian gene expression panel. This suggests a particular role for the circadian clock’s genetic architecture in driving drug responses to topotecan.

In summary, our analysis underscores the circadian clock’s critical role in determining drug sensitivity across a wide range of drugs. We demonstrate that a combination of circadian features, rather than a single factor, discriminates between high and low drug responses. Through the introduction of the “chronosensitivity index”, we provide a robust metric for quantifying the circadian clock’s impact on differentiating between high and low drug sensitivity groups, thereby identifying potential drugs that act in a circadian-dependent manner.

## Discussion

Despite the wide acknowledgement of the fundamental role that the circadian clock occupies in the progression of diseases and in the modulation of treatment responses (Lévi *et al*, 2010; Mormont & Levi, 2003; Sulli *et al*., 2019), our understanding of the underlying molecular mechanisms remains fragmented. Using a high-resolution circadian phenotyping approach to thoroughly characterize circadian rhythms in cancer models, we discovered strong circadian rhythms in numerous breast cancer cell models. Among these, models of the most aggressive TNBC subtype showed distinct circadian rhythms, challenging the general expectations that the circadian clock is rather dysregulated in highly transformed cancer (Mormont & Levi, 2003; Ye *et al*., 2018) (**Fig. 2**). Our measurements of the circadian clock in the non-malignant epithelial cell line MCF10A, luminal A (LumA) breast cancer cell line MCF7, and the osteosarcoma U-2 OS cells aligns with prior research (Baggs *et al*., 2009; Börding *et al*, 2019; Lellupitiyage Don *et al*., 2019; Lin *et al*, 2019), proving the applicability and flexibility of our approach for deep circadian phenotyping across tissue and cancer types.

Recent work revealed subtype-dependent clock functionality in breast cancer and that breast cancer clocks are critically regulated by estrogen responsiveness (Li *et al*., 2024). Our analysis of multiple cell models per breast cancer subtype and our broad set of circadian parameters confirms variations in clock strength across breast cancer subtypes, however, we also revealed substantial variability within the individual subtypes of breast cancer. This variety of distinct clock features motivated us to define circadian clock-based phenotypes within the panel of cell lines tested. In detail, we systematically categorized cell models into groups with functional, weak, unstable, or dysfunctional clocks, based on representative parameters indicative of circadian strength, stability, and periodicity (**Fig. 3C**). This classification diverged from traditional subtyping, exposing functional clocks in only two of the three tested estrogen-positive LumA cell models (Sun *et al*, 2014), alongside three out of four estrogen-negative TNBC models. Together, the defined clock phenotypes offer a complementary viewpoint on the subtyping of breast cancer which could refine the assessment of cancer models for circadian-aligned therapeutic strategies and improve the prediction of these therapeutic outcomes. The previously mentioned study on assessing global rhythm strength in breast cancer patient samples following an informatic approach emphasizes the importance of tumor subtype-specific analysis of circadian rhythms (Li *et al*., 2024). Our findings further support this need and highlight the importance of assessments that consider the distinct characteristics of each tumor individually.

On the molecular level, circadian clocks ensure coordinated rhythmicity from the individual cell to whole organism level by organized TFFLs, regulated through the interaction of multiple circadian clock genes (Chaix *et al*., 2016; Takahashi, 2017). The functionality of the circadian clock is highly dependent on genomic integrity, as proved by animal studies revealing distinct phenotypic effects for mutations of mammalian clock genes (Ko & Takahashi, 2006). Examining the potential of inferring our established circadian-based phenotypes from publicly available genetic profiles of the cell models, we found high mutation burdens in circadian genes and compromised clock functionality, particularly within the unstable and weak phenotypes which underscores the genetic basis for circadian disruption in these groups. In contrast, cell models with functional clocks exhibited lower mutation load and less severe genetic alterations, suggesting higher genomic stability in the circadian clock mechanism (**Fig. 3E**).

Illuminating the link between the circadian clock and drug sensitivity, we found strong correlations between selected circadian features and drug sensitivity metrics (**Fig. S5A**). Considering absolute correlation values per circadian metric, we revealed the circadian amplitude as most correlated metric with the maximal drug responses and the concentration at half-maximal drug growth inhibition (*GR*_50_) for 9 distinct drugs. However, despite these significant correlations, the overall correlations between individual circadian clock features and drug sensitivity metrics were rather moderate. This suggests that drug sensitivity is not shaped by a single circadian factor, but rather by the collective influence of multiple circadian features which we confirmed by linear discriminant analysis. Furthermore, we found that the most distinctive circadian features vary depending on the specific drug or sensitivity parameter in question. For example, while strength-related circadian parameters such as the rhythmicity index and circadian amplitude were strong predictors for the drug sensitivity to the DDR-inducer cisplatin (**Fig. 4D**), parameters indicative for clock stability were mostly contributing to the discrimination into sensitivity groups to the HSP90 inhibitor 17-AAG (**Fig. 4F**). These results suggest a complex relationship between circadian rhythms and drug responses that needs to be evaluated on a drug-by-drug basis.

With the “chronosensitivity index” we systematically quantified the dependency of drug responses on the circadian clock, based on the ability of circadian parameters or circadian gene expression levels to effectively classify cell models into distinct sensitivity groups. By that we identified circadian clock-sensitivity relationships for most drugs studied, highlighting a global influence of the circadian system on pharmacodynamics (**Fig. 4H** and **I**). Importantly, 17-AAG was the highest-ranking drug in chronosensitivity index suggesting a particularly reliable separation into high or low drug response groups by circadian clock parameters for the cell line panel tested. In an earlier study which performed a high-throughput screening of time-of-day effects on anticancer drug activity, Lee et al. identified high time-of-day-specific action for the HSP90 inhibitor 17-AAG, mediated by the circadian regulation of the cell cycle (Lee *et al*., 2021). Our findings complement these findings, underscoring the critical influence of circadian rhythms on drug efficacy, highlighting the importance of integrating circadian biology into pharmacological research and treatment strategies. Though, while circadian parameters and gene expression profiles proved to offer insights into drug responses, differences in their predictive power for certain drugs, such as topotecan, suggest areas for future research.

## Material and Methods

### Experimental methods Cell lines and cell culture

The TNBC cell lines HCC1143, HCC1806, HCC1937, HCC38, HCC70, and MDAMB468 were acquired from the American Type Culture Collection. BT549, CAL51, HCC1428, MDAMB231, MDAMB436 and SUM149PT cells were kindly gifted by the Sorger lab (Harvard Medical School, Ludwig Cancer Center, Boston, USA). MCF10A, MCF7, and T47D cells were kindly provided by the Brugge lab (Harvard Medical School, Ludwig Cancer Center, Boston, USA). The U-2 OS reporter cell lines originated from the Kramer lab (Charité, Institute for Medical Immunology, Berlin, Germany). MCF10A cells were cultured following the Brugge lab’s media recipe including a DMEM/F12 medium base (Gibco) supplemented with 5% horse serum (Gibco), 20 ng/ml EGF (Peprotech), 0.5 mg/ml Hydrocortisone (Sigma), 100 ng/ml Cholera Toxin (Sigma), 10 µg/ml Insulin (Sigma) and 1% penicillin-streptomycin (Pen-Strep, Gibco). All other cell lines were maintained in RPMI-1640 medium (Gibco) supplemented with 10% fetal bovine serum (FBS) (Gibco) and 1% Pen-Strep. Bioluminescence recordings and live imaging required phenol red-free FluoroBrite DMEM medium, supplemented with 10% FBS, 300 mg L-Glutamine (Gibco) and 1% Pen-Strep. Cells were cultured in a controlled 37 °C and 5% CO_2_ environment and routinely tested for mycoplasma for quality assurance.

### Generation of luciferase and fluorescence reporter cell lines

HEK293T cells at 80% confluency were retained in RMPI-1640 medium supplemented with 10 mM HEPES and transfected with a mix of 8.4 µg lentiviral expression plasmid (pAB-mBmal1:Luc-Puro or plenti6-mPer2:Luc-Blas), 6 µg psPAX2 (Addgene #12260) and 3.6 µg pMD2G (Addgene #12259) to produce lentivirus carrying circadian luciferase reporters. Similarly, lentivirus for the red-fluorescent nuclear reporters were produced using a mix of 1.8 μg gag/pol packaging plasmid (Addgene #14887), 0.7 μg pRev packaging plasmid (Addgene #12253), 0.3 μg VSV-G envelope plasmid (Addgene #14888) and 3.2 μg of EF1α-mKate2-NLS plasmid. The transfections were performed using Lipofectamine 3000 (Invitrogen) according to the manufacturer’s instructions. Virus was harvested and filtered through a 0.45 μm filters (Millipore) 48 h and 72 hours post-transfection. For transduction, target cells at 70% confluence underwent a 6-hour incubation with a mix of 1 ml lentivirus-containing supernatant, 8 μg/ml protamine sulfate (Sigma) and 10 μM HEPES (Gibco). Post-incubation, cells were washed with PBS (Gibco) and maintained in standard culture medium. After 2 days, antibiotic selection of transduced cells was initiated using either 5 μg/ml of blasticidin (Adooq) or 2 μg/ml of puromycin (Gibco), depending on the resistance marker present in the lentiviral expression vector. Details on how the different U-2 OS knockout cell lines were developed are described in the original publication (Börding *et al*., 2019).

### Circadian bioluminescence recordings

Cells expressing luciferase reporters under the control of the *Bmal1* or *Per2* promoters were plated in 35-mm dishes (Nunc) to reach approximate confluence by the next day. To synchronize single-cell circadian clocks in cell populations, a standard dose of 1 µM dexamethasone (Balsalobre *et al*, 2000; Finger *et al*., 2021) (Sigma, dissolved in EtOH) was applied. Following a 30-minute incubation period, cells were rinsed once with PBS before adding imaging medium containing 250 μM D-Luciferin (Abmole). To prevent the imaging medium from evaporating during the duration of the bioluminescence measurements, the dishes were sealed with parafilm (Finger *et al*., 2021). Bioluminescence was then recorded every 10 minutes for a span of 5.7 days, utilizing an incubator-embedded luminometer (LumiCycle, Actimetrics).

### Live-cell imaging

Continuous live-cell monitoring was performed utilizing an incubator-embedded live-cell widefield microscope (Incycte, Essen BioScience). Cells expressing the fluorescent nuclear marker EF1α-mKate2-NLS were plated in 48-well plates (Falcon) at a seeding density where cell growth saturated under normal growth conditions by the conclusion of each experiment. Growth was measured in both brightfield and red-fluorescent channels with an excitation between 567–607 nm and emission between 622–704 nm. The acquired images were subjected to per-frame analysis for nuclei counting using the integrated software of the Incucyte system, followed by data processing in MATLAB. In experiments involving pharmacological treatments, cell seeding was done 1 day prior to the initiation of imaging. We employed a 4x objective to take images across two fields of view for each well, at intervals of 1–2 hours, over a period of 4 days. These experiments were replicated across two separate plates for each condition. In the case of cisplatin treatment studies, we conducted the experiment in a single plate per condition, capturing images across nine fields of view with a 10x objective. For the assessment of unperturbed growth dynamics, we seeded cells at approximately 10% confluency, two days before initiating the live imaging. Using a 10x lens, we obtained nine snapshots per well at 1–2-hour intervals, for four days.

### Drug perturbations

Stock solutions for drugs were prepared in DMSO and stored at -20°C, except for cisplatin (Sigma) that was prepared in 0.9% NaCl and kept at room temperature. For dose-response experiments, drugs were freshly diluted in a logarithmic series (5–6 points, log4) using their respective solvents just before application. The following concentration ranges were evaluated: 5-fluorouracil (5-FU, Sigma), alpelisib (Biozol), and olaparib (Adooq) were tested from 100 to 0.1 µM; torin2 (Sigma) and alisertib (Hölzel) from 10 to 0.01 µM; adavosertib (Biocat) from 10 to 0.04 µM; doxorubicin (Hölzel) from 1 to 0.004 µM; and paclitaxel (Hölzel) from 0.4 to 0.0004 µM. Cisplatin underwent a 10-point log2 dilution series, spanning 70 to 0.14 µM. The compounds were mixed media to achieve a final volume of 9% of the well and administered to the cells 24 hours post-seeding. Cisplatin doses were given 48 hours post-seeding. Control groups received the respective solvent to account for potential solvent-based effects on growth. Growth of the cells was tracked by long-term live-cell imaging over a minimum period of 4 days, as previously detailed.

### Computational methods

#### Time-series analysis of circadian signals

##### Signal pre-processing

An initial dexamethasone-related signal peak within the first 5 hours was uniformly removed from all raw time-series. To standardize the duration of recordings, time-series exceeding 137.7 hours (∼5.7 days) were shortened accordingly. In the case of the MDAMB468 *Per2*-Luc reporter cell line, one replicate required a further adjustment by shortening the dataset by 8 hours (129.7 hours). This was due to an anomalous, intense, and brief peak during the last recording hours that skewed the analysis of circadian parameters, and that was not present in other replicates. The signals were detrended with a 48-h cut-off period using a sinc filter. Signal normalization was achieved by inverting the continuous amplitude envelope of the detrended signal (Mönke *et al*, 2020). The amplitude envelope was calculated by continuous wavelet transform with a time window of 48 hours. The signal pre-processing steps were performed in the open-source software package pyBOAT (Mönke *et al*., 2020) (version 0.9.12), accessed via the Anaconda Navigator (version 2.5.0).

##### Autocorrelation analysis

Autocorrelation of detrended signals was calculated using the ‘autocorr’ MATLAB function. Rhythmicity strength was determined from autocorrelation values at the second peak in the correlogram using the ‘findpeaks’ MATLAB function, with the corresponding abscissa (lag) reflecting the main period. To facilitate peak detection and to filter out non-periodic samples from further analysis, autocorrelation values were smoothed using a gaussian filter and peaks outside the 95.5% confidence intervals or below 0 were not considered for further analysis.

##### Multiresolution analysis

Decomposition of detrended signals into the underlying frequency bands, referred to as wavelet details *D*_j_, was done by a discrete wavelet transform based multiresolution analysis (Leise & Harrington, 2011), ensued by a final smooth. The analysis was carried out in accordance with the method outlined by Myung, Schmal and Hong *et al* (Myung *et al*, 2018), employing the ‘PyWavelets’ Python library and utilizing a *db20* wavelet from the Daubechies family for the transformation. For each wavelet detail *D*_j_, which encapsulates a frequency span from 2^j^Δt and 2^j+1^Δt (where j=1, 2, 3, …), the time series was resampled by reducing the sampling interval from Δt=10 min to Δt=30 to capture a circadian frequency range between 16–32 hours (Leise, 2013).

##### Continuous wavelet transform

Continuous wavelet transform (CWT) was employed for the identification of a time-series signal’s main oscillatory feature, referred to as ridge, across the entire recording time. The application of this wavelet-based spectral analysis varied with the specific readout parameter being examined. To increase the detectability of oscillatory patterns associated with the “period” and “phase” readout parameters, the CWT was applied to detrended amplitude-normalized signals. Furthermore, an adaptive ridge detection threshold was set according to the quarter-maximal spectral power of each signal (**Fig. S1A**, right panel). Conversely, for the readout parameters “amplitude” and “ridge length”, indicative of circadian strength and robustness, the analysis was conducted on detrended, unnormalized signals with a global ridge detection threshold that results in a broad distribution of ridge lengths across all samples tested and which we set to a quarter of the median half-maximal wavelet power from the aggregate of samples (**Fig. S1aA**, left panel). Except for the “ridge length” parameter, we focused on analyzing ridges detectable for a minimum duration of 48 hours, thereby ensuring the tracking of at least two full circadian cycles for our subsequent analysis. The CWT was performed with pyBOAT using the Python programming language in the Spyder environment (version 5.4.5). The instantaneous phase differences between *Bmal1* and *Per2* was determined using the ‘atan2’ function, and for visual representation in polar histograms, the ‘polarhistogram’ function from MATLAB.

##### Weighting of CWT parameters

To account for variations in ridge lengths from which CWT period, amplitude, and phase parameters were derived, we weighted the importance of each sample through a sigmoidal fitting method applied to the lengths of the respective ridges (**Fig. S1B**), based on the formula:

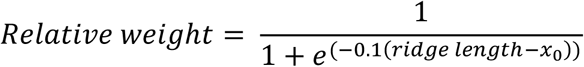

Here, the steepness of the curve was uniformly set to -0.1, while the reflection point of the curve, *x*_0_, lied between the highest and lowest ridge length. This method ensured that parameters from longer ridge lengths had a higher relative weight for median and mean calculations, as opposed to lower-weighted parameters from shorter ridge lengths.

##### Amplitude envelope decay rate calculation

Decay rates of amplitude envelopes used for later correlation analyses were derived by applying an exponential decay model to detrended signal amplitudes:

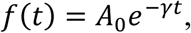

where *A*_0_ denotes the initial amplitude, *t* the time, and *γ* the decay rate. Only decay rates from fits where *R*^2^ ≥ 0.6 were considered for further analysis. For the analysis of circadian coupling strengths, the model was extended to fit signals to an exponentially decaying sinusoidal oscillation.

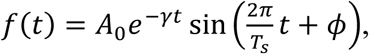

where *T_s_* is the oscillation period, and *ɸ* adjusts for phase shifts. Again, values obtained from fits where *R*^2^ ≥ 0.6 were used for further analysis.

##### Simulating circadian population signals

The intercellular coupling was modeled assuming cells as mean-field coupled Poincaré oscillators. These dynamics were described in Cartesian coordinates as:

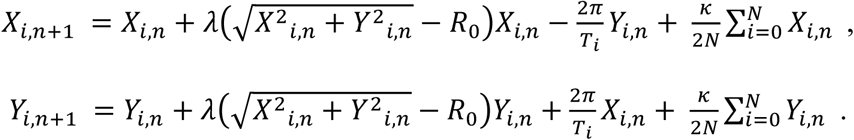

Here, *n* represents the number of identical oscillators, *κ* the circadian coupling strength, and *T_i_* the inherent period. To simulate the model we used the Euler’s method with a timestep of Δt=0.1. For simplicity we set *R*_0_=*λ*=1, as these parameters had no significant impact on the outcome of the experiment. To reenact the in vitro resetting, all oscillators were given similar initial phases by Uniform(0, π/5), and periods given by a normal distribution Norm(24, 3). Further, we set *N* to 300 to simulate enough oscillators and to avoid edge case results while keeping the simulation relatively small. Identical oscillators and uniform coupling in the simulations were justified by our assumption of absent spatial correlations in the experiment. The cumulative cell signal was modeled by summing individual oscillator outputs 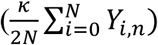, varying *κ* ∈ [0.00001 : 0.1], with amplitudes scaled by 0.5 for compatibility, while keeping a constant seed to ensure similar distribution of periods for all simulations.

##### Statistical analysis

Significance in variations in circadian parameters between knockout and wild-type U-2 OS cells were assessed using a two-sample *t*-test through the M ATLAB ‘ttest2’ function. Variations were considered significant with corresponding *p*-values ≤ 0.05.

### Analysis of growth dynamics

Growth data acquired by long-term live-cell imaging was smoothed by robust local regression of weighted linear least squares combined with a 2^nd^ degree polynomial model, as implemented by the ‘rloess’ function MATLAB. The exponential growth rate, denoted as *k*, for each time unit *t* was determined by normalizing the cell counts to their initial values at timepoint zero (*y*_0_) and applying an exponential model to the 4-day growth trajectories:

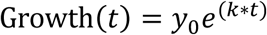

For the BT549 and MDAMB436 cell lines fluorescent reporter lines were not available, which is why growth was quantified based on confluency levels. For all other cell models, nuclear counting was employed to calculate growth rates. Fitting of the exponential model was employed by the ‘fit’ function in MATLAB.

### Circadian parameter-based phenotype analysis

The relationship across different circadian parameters and growth rates were computed and interpreted using the Pearson correlation coefficient and corresponding *p*-values on combined *Bmal1*-*Per2* data, averaged across breast cancer cell lines. From the reduced set of eight circadian parameters and the growth rate, unsupervised dimensionality reduction by principal component analysis (PCA) was executed using the ‘PCA’ function of the ‘sklearn decomposition’ Python language module with the exact full singular value decomposition solver and two output components. Min-max scaling of the circadian parameters was applied prior to processing. The reduced 2-dimensional space of *Bmal1*-*Per2* circadian features of each cell line was plotted in a biplot together with each feature’s loading by annotated arrows, revealing three distinct clusters. The resulting component loadings were plotted separately in bars to facilitate the interpretation of each circadian feature’s contribution. Based on the feature contributions, one representative parameter was selected from each of the three PCA clusters (period, phase difference variability, circadian component) and their min-max scaled values were plotted in a three-dimensional space. Discretization using the *k*-means algorithm and *k*=4 revealed the four clock phenotype clusters functional, unstable, weak, and dysfunctional (within cluster sum of squares: 0.79, cluster silhouette score: 0.45).

### Circadian gene expression and mutation analysis

Gene expression data and somatic mutation information of circadian clock genes were sourced from the Cancer Cell Line Encyclopedia Dependency Map (CCLE DepMap, available at https://sites.broadinstitute.org/ccle/datasets, gene expression: 2022-Q2, mutation profiles: 2023-Q2) (Barretina *et al*., 2012). Cell models were grouped together based on their expression values of core circadian clock genes using the ‘seaborn’ library’s ‘clustermap’ function, utilizing Euclidean distance for measurement and the complete linkage approach for clustering. We extended our mutation analysis from the set of 16 core clock genes to additional 44 circadian clock associated genes (see **Supplementary Table 1** for gene lists). Cell lines were hierarchically clustered by their mutational burden ranging from zero to greater or equal two, using the hamming distance metric and single linkage method. Depending on the severity of the mutation, each mutation was assigned a value between 1 to 3 (with 3 reflecting damaging mutations), and final mutation scores per clock phenotype or classical subtype were calculated from averaging individual mutation scores across cell line models.

### Circadian feature and gene expression mapping

Min-max scaled gene expression values of sixteen core clock genes across TNBC cell line models were clustered using the ‘clustermap’ function of the ‘seaborn’ Python language module with the Euclidean distance metric and complete linkage method. The Uniform Manifold Approximation and Projection (UMAP) algorithm of the same Python language module was applied on both, the circadian parameters, and circadian gene expression values, to project TNBC cancer cell lines in a two-dimensional space. The number of nearest neighbours selected to construct the initial high-dimensional graph was set to three.

### Estimation of growth rate inhibition values

Drug response data acquired by long-term live-cell imaging was smoothed using the same approach as described for the growth analysis. Following the method outlined by Hafner *et al*. (2016), the growth rate inhibition (*GR*) at a given time *t* and for each concentration *c* was calculated using:

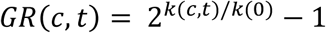

where *k*(*c*, *t*) represents the growth rate with drug treatment, and *k*(0) signifies the growth rate of untreated control cells. To obtain final drug sensitivity values, dose-dependent *GR* values were modeled against a sigmoidal curve, expressed as:

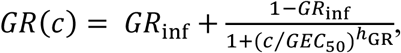

with the parameters defined as described in the original publication (Hafner *et al*., 2016). Fitting of the sigmoidal curve was executed with the MATLAB ‘fit’ function.

### Correlation and linear regression analysis

Pearson’s linear correlation coefficients between drug sensitivity metrics and circadian parameters were calculated and clustered using the ‘corr’ and ‘clustergram’ functions in MATLAB, respectively. For clustering, the ‘correlation’ distance metric was applied. Corresponding linear regression analysis was employed by the MATLAB ‘fitlm’ function.

### Predicting drug sensitivity from circadian clock features

Drug sensitivity prediction analysis was performed using the supervised linear discriminant analysis algorithm by the ‘sklearn discriminant_analysis’ Python language module. The default parameters were retained, using the exact full singular value decomposition solver and a single component (LD1). We applied a median-based binary approach to categorize cell models into high and low drug sensitivity groups for cisplatin (own dataset) and 24 additional drugs that we obtained from the CCLE pharmacological profiling data set archive (2015-Q1) (Balsalobre *et al*., 2000). These sensitivity groups served as target variable for the discrimination analysis using either the circadian parameter set (combined *Bmal1*-*Per2* data), or the CCLE core clock gene expression values. The resulting linear discriminant vector was jittered along the y-axis to avoid overlapping of the data points in the 1-dimensional space. Individual contributions to the obtained discriminative information of the parameters were plotted alongside each linear discriminant plot, and combined sum up to 100%. The discrimination performance (“chronosensitivity index”) was calculated from the BCD-WCD ratio, where the between-cluster-distance (BCD) is the Euclidean distance between the mean LD1 values of each class, and the within-cluster-distance (WCD) is the average Euclidean distance of class members to their respective class mean. The minimum BCD-WCD ratio to deem effective discrimination in a single dimension is equal to two, with BCD=2*WCD. This ensures that the centers of both clusters are at least twice as far away as the distance of each member to its respective class mean.

## Data availability

The experimental time series data and data tables for all results of this study are available upon request.

## Code availability

Code used for the data analysis in this work is available upon request.

## Abbreviations

5-FU: 5-fluorouracil
AC: autocorrelation
ATCC: American type culture collection
BCD: between-cluster-distance
BL1: basal-like1
BL2: basal-like2
BMAL1: brain and muscle arnt-like protein-1
CI: confidence interval
CLOCK: circadian locomotor output cycles protein kaput
CRY: cryptochrome
CWT: continuous wavelet transform
dKO: double-knockout
DMEM: Dulbecco’s Modified Eagle Medium
DMSO: dimethyl sulfoxide
DWT: discrete wavelet transform
EGF: epidermal growth factor
GR: growth rate inhibition
HSP90: heat shock protein 90
LDA: linear discriminant analysis
Luc: luciferase
MES: mesenchymal
mTOR: mammalian target of rapamycin
NLS: nuclear localization sequence
PARP: poly ADP ribose polymerase;
PCA: principal component analysis
PER2: period2
PI3K: phosphatidylinositol 3-kinase/alpha serine/ threonine-protein kinase
ROR: RAR-related orphan receptor
RPMI: Roswell Park Memorial Institute medium
SCN: suprachiasmatic nucleus
sKO: single-knockout
TTFL: transcriptional-translational feedback loop
TNBC: triple-negative breast cancer
TPM: transcripts per million
UMAP: uniform manifold approximation and projection
WCD: within-cluster-distance.

## Acknowledgements

We would like to express our thankfulness to Nica Gutu for kickstarting the modelling ideas. We also thank Anna-Marie Finger and Astrid Grudziecki for their guidance in the bioluminescence recordings as well as Francesca Müller-Marquardt for her assistance in the recordings. Lastly, we thank the laboratories of Ingeborg Tinhofer-Keilholz and Ulrich Keilholz for their continuous feedback on the project. The results are part of a project funded by the German Federal Ministry of Education and Research (BMBF) through the e:Med Juniorverbund DeepLTNBC TP3-01ZX1917C. C.E. was partially supported by the Deutsche Forschungsgemeinschaft (DFG, German Research Foundation)–RTG2424/CompCancer – project number: 377984878. M.S.N. acknowledges support from the Novo Nordisk Foundation (NNF20OC0064978). C.S. acknowledges support from the DFG–SCHM 3362/4–1 project number: 511886499.

---

**Box 1.**
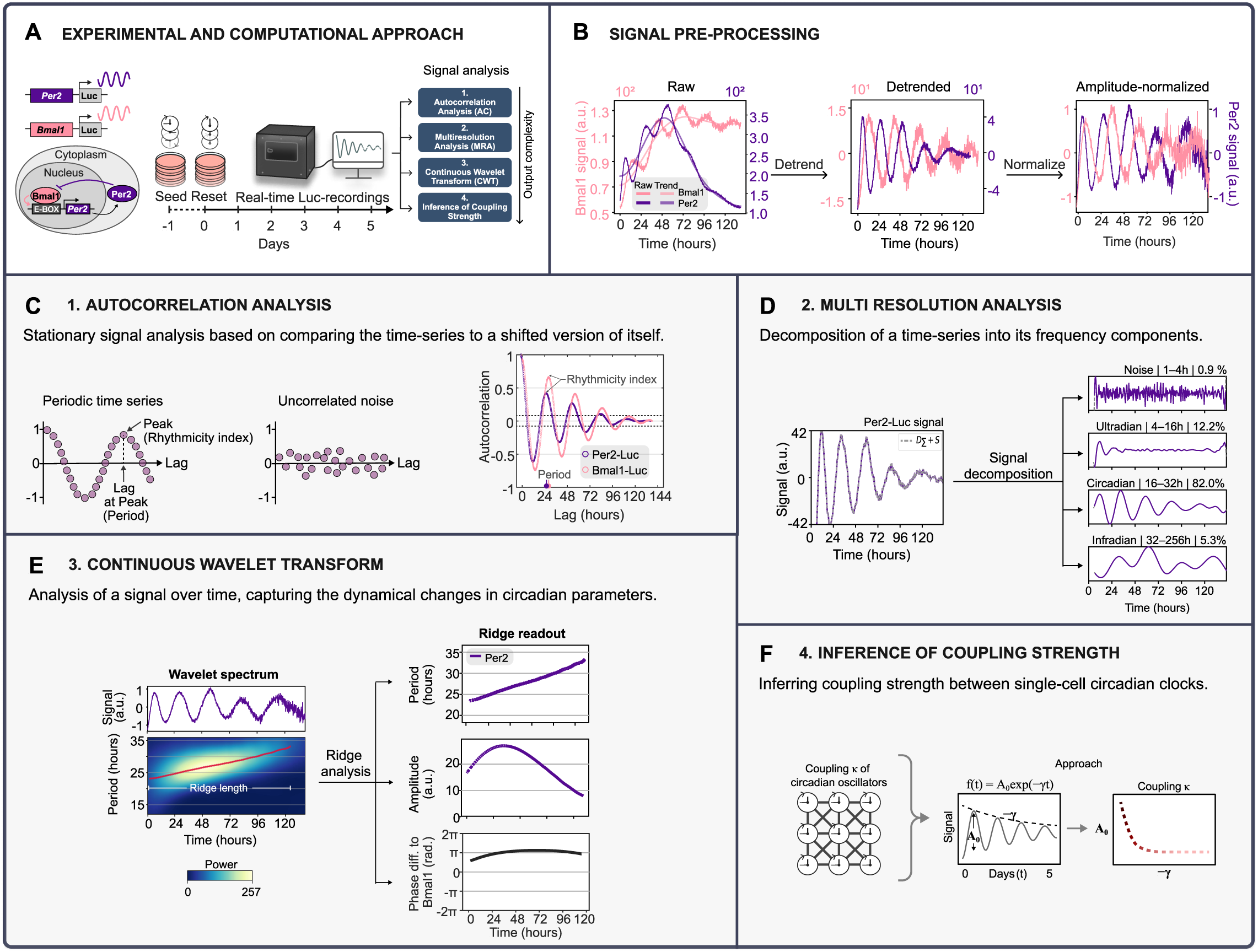
**(A)** Schematic of the experimental and computational approach to deep phenotype circadian rhythms. Stable circadian reporter cell lines expressing Luciferase (Luc) reporters for either *Bmal1* or *Per2,* two core clock genes of the circadian clock network, were generated by lentiviral transduction. The circadian clocks of individual cells in a cell population were reset, and signals from the reporters were monitored in real-time by biolumenscence recordings spanning multiple days. **(B)** Raw signal (left plot) and pre-processed signal traces (middle and right plots) of the TNBC cell line HCC1143 Bma/7-Luc and Per2-Luc. Pre-processing of raw signals involved detrending (middle plot) and, for parts of the analysis, amplitude-normalization (left plot). **(C)** Autocorrelation analysis (AC) of detrended HCC1143 Bma/7-Luc and Per2-Luc signals. The arrow indicates the rhythmicity index and corresponding period of the time-series (lag). Dashed lines=95.4% CI. **(D)** Multiresolution analysis of detrended HCC1143 Per2-Luc signal to decompose the signal into four frequency components, %=fraction of each component to the total signal. (E) Continuous wavelet transform analysis on detrended and amplitude-normalized HCC1143 Per2-Luc signal (top left plot), showing time-resolved signal periods in a wavelet spectrum (bottom left plot). The red line marks the main oscillatory component (ridge). The right panel shows the corresponding ridge readout for the period (top), amplitude (middle), and phase difference to a respective HCC1143 Bma/7-Luc signal (bottom). (F) Approach to calculate the coupling strength (k) of circadian oscillators within a population of cells from the signal’s amplitude (A_o_) and exponential decay rate (*γ*).

**Figure S1.**
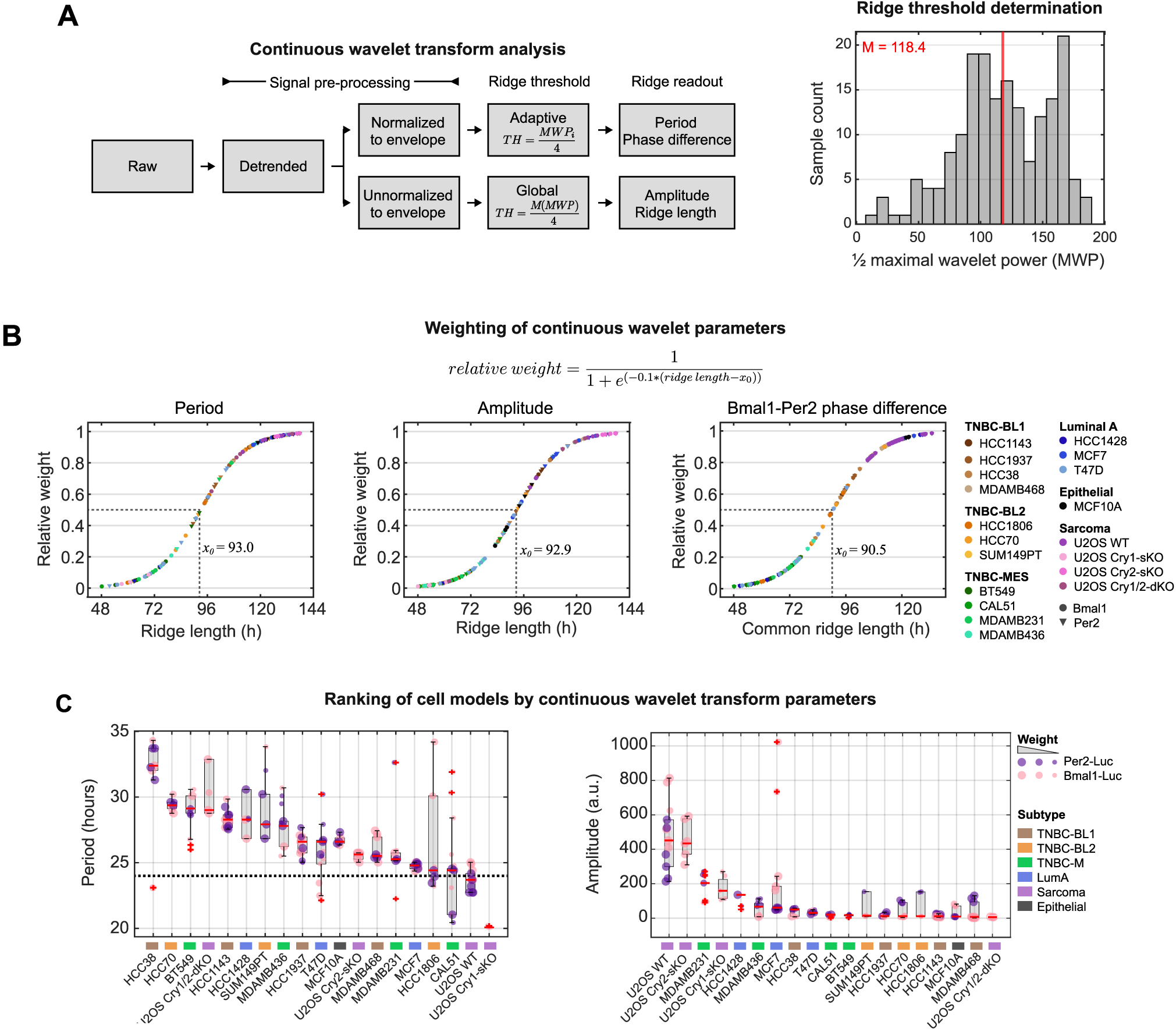
Continuous wavelet transform analysis. (**A**) Approach to extract ridge readout parameters (left panel). Ridge thresholds (TH) applied to the power spectrum were either adaptive, set to a quarter of the sample’s maximum wavelet power (MWP), or global, set to a quarter of the median of the maximum wavelet power of all samples combined (right panel, n=180 samples). **(B)** Calculation of relative weights of continuous wavelet transform (CWT) parameters through sigmoidal fitting of ridge lengths underlying the extracted periods and amplitudes (left and middle panel) or of the common ridge lengths of *Bma/1-* and Per2-phases for the calculation of phase differences (right panel). Cell line models are color-coded by their subtype. The hill coefficient of the sigmoidal model was set to -0.1 and calculated inflection points (x0) of the fitted curve are marked in the plots. **(C)** Weighted boxplot of median periods (left panel) and amplitudes (right panel) from *Bma/1-* and Per2-signals of the specified cell models, with the bottom and top edges of the boxes representing the 25^th^ and 75^th^ percentiles, respectively. Extending whiskers represent data points within 1.5 times the interquartile range from lower and upper quartile. Relative weights of the samples, calculated as shown in B, are indicated by datapoint sizes: small (:S 0.33), intermediate (> 0.33 and :s 0.66), and large (> 0.66). Red horizontal lines denote median values, red crosses mark outliers.

**Figure S2.**
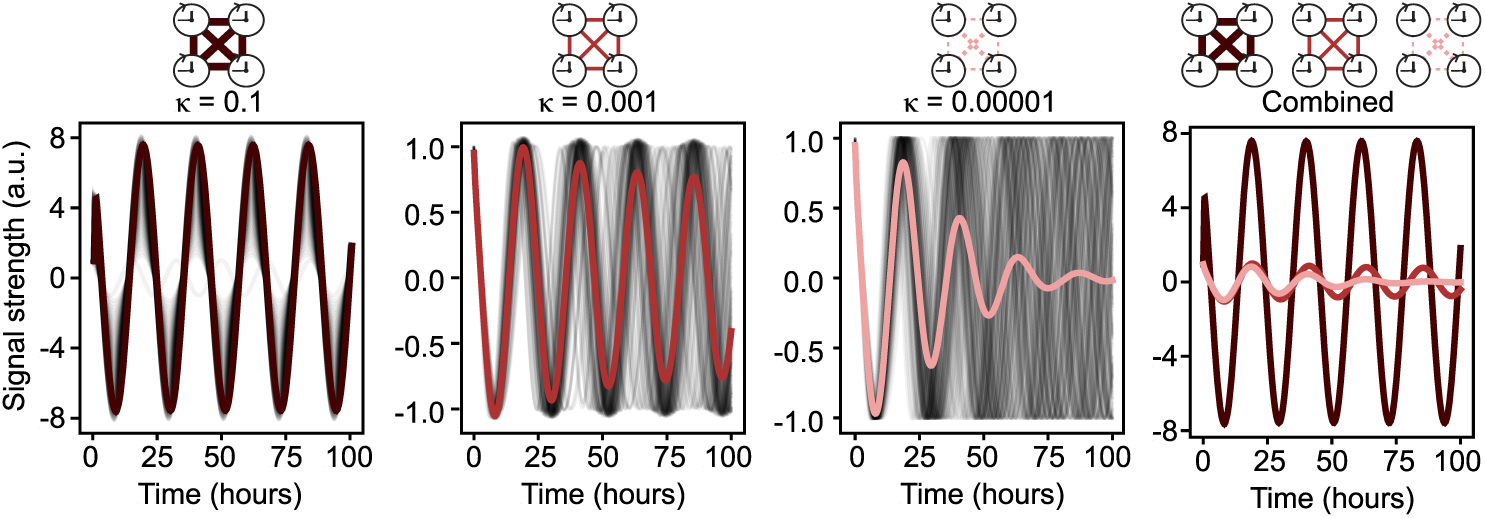
Simulation of coupled circadian oscillators. Simulated oscillators of varying coupling strengths κ. Individual traces of each oscillator are shown in black, collective signals are shown as thick lines with gradients of red, based on their coupling strength, where dark red denotes high coupling strength, and pink refers to low coupling. The right panel is a composite of all collective signals.

**Figure S3.**
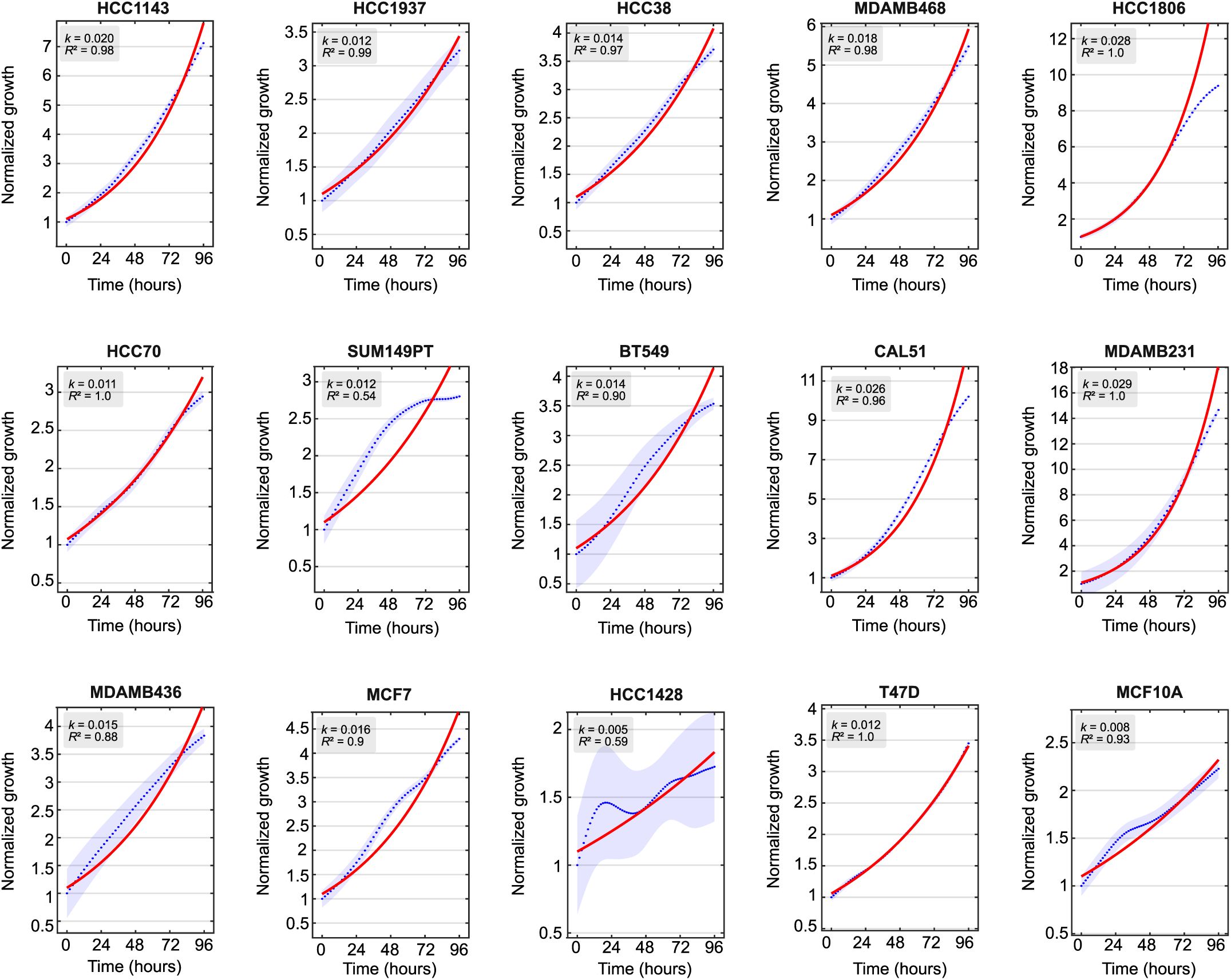
Capturing growth dynamics by long-term live cell imaging. Normalized growth curves (blue dots) and exponential fits (red lines) for the indicated cell line models, yielding growth rates (k), and fit accuracies (R^2^). Growth assessment was based on nucleus counts, except for BT549 and MDAMB436, where confluency was used. Data represents the mean±s.d. of 9 images taken in a single well, with the shaded area representing the standard deviation across the images.

**Figure S4.**
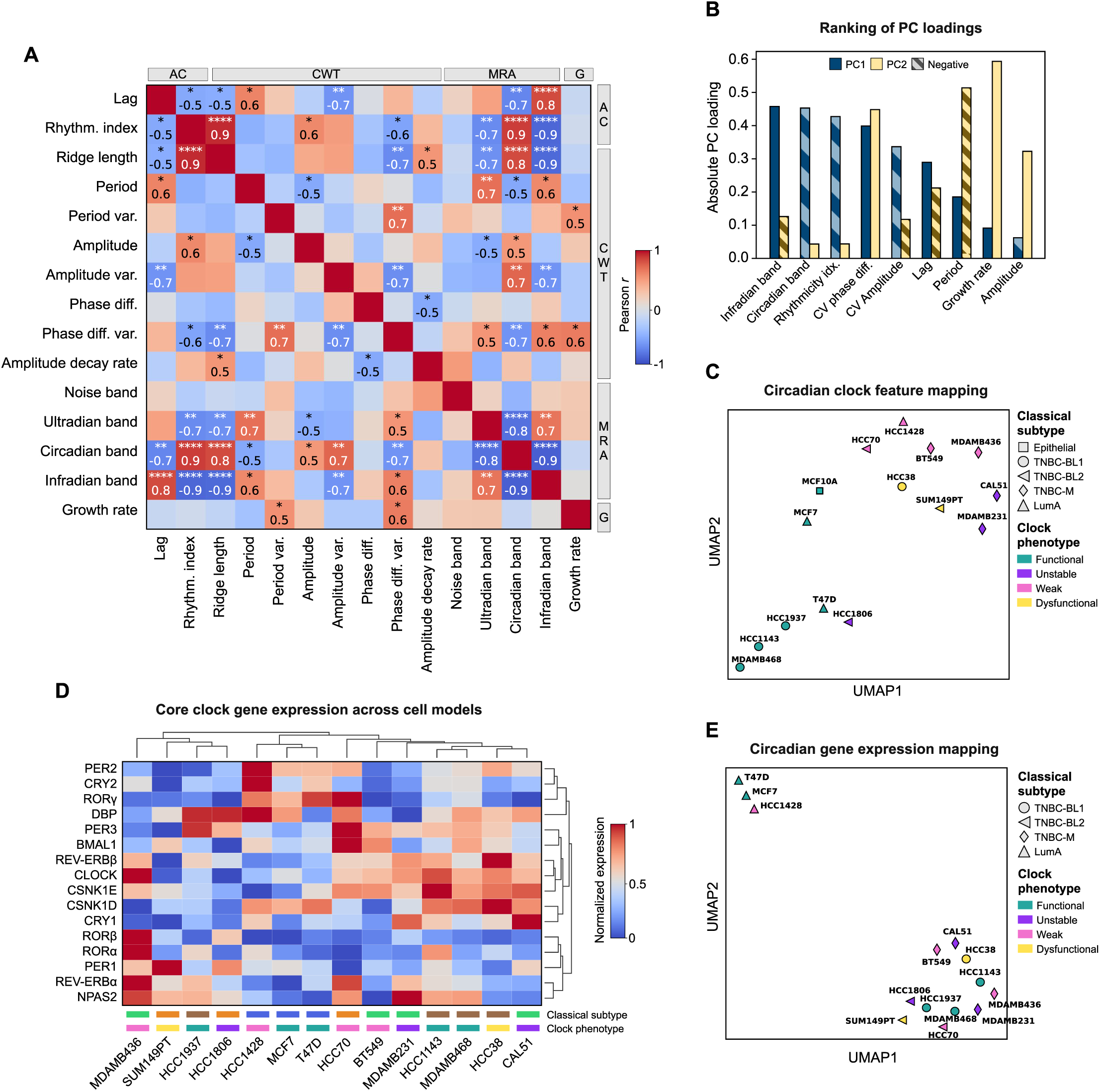
Mapping circadian clock features and gene expression. **(A)** Pearson correlation coefficients between the complete set of *Bmal1-Per2-* averaged circadian features and growth rates across all breast cancer cell models and the epithelial MCF10A cell line (n=15 cell lines). Parameters are categorized by their approach of calculation (refer to Figure 2a for full names; G=growth). Displayed are statistically significant correlation values, where *, **, ***, and **** indicate p-values ≤ 0.05, 0.01, 0.001, and 0.0001, respectively. (B) Ranking of absolute PC loadings for each circadian and growth parameter, corresponding to the PCA biplot shown in Figure 3B. Parameters are sorted in descending order based on their absolute contribution in the first principal component. Negative values are shaded. (C) Uniform Manifold Approximation and Projection (UMAP) of cell models based on selected circadian clock features shown in B. The classical subtype and clock phenotype of each model is illustrated by different markers and colors, respectively. Nearest neighbours=3. (D) Cluster map of core clock gene expression values across breast cancer cell models, using the Euclidian distance method. Color-coded rectangles above the x-labels indicate the classical subtype and clock phenotype. Refer to Figure 3B and 3C for color-coding. (E) UMAP of cell models based on core circadian gene expression values shown in D. The classical subtype and clock phenotype of each model is illustrated by different markers and colors, respectively. Nearest neighbours=3.

**Figure S5.**
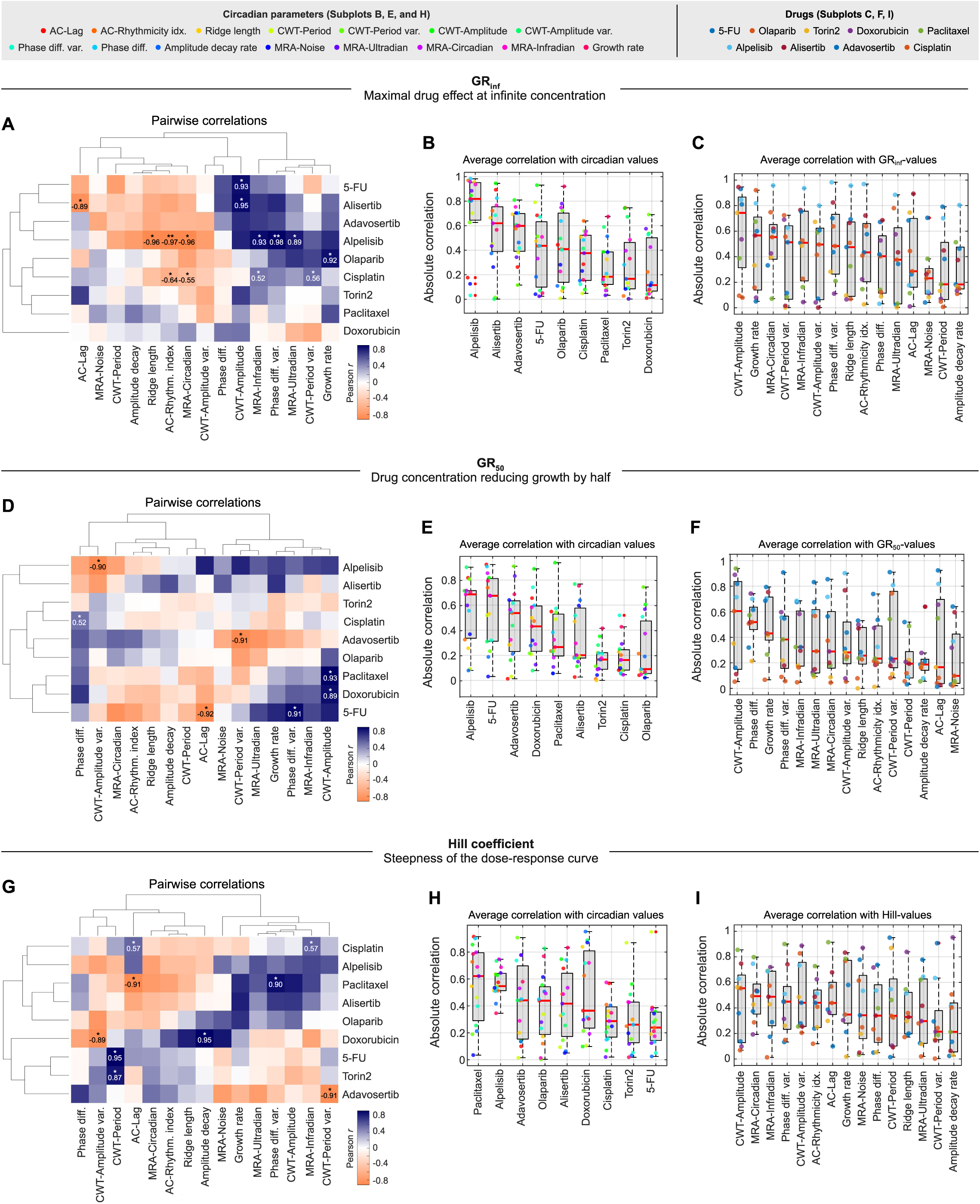
Relationship between circadian clock features and drug sensitivity metrics to various drugs, **(A)** Hierarchical clustering of Pearson correlation coefficients between GR;nr values for 9 drugs (rows) and 15 circadian clock and growth parameters (columns, averaged *Bma/1-Per2* data). Shown are statistically significant correlation values, where *, and **, indicate p-values ≤ 0.05, and 0.01, respectively. n=15 cell lines. (Band **C)** Ranking of the absolute correlation between GR;nr values of various drugs and circadian clock/growth parameters, accumulated either by drug **(B,** n=15 parameters), or by parameter **(C,** *n=9* drugs). Data points are color-coded according to the key presented at the top of the page. Bottom and top edges of the boxes represent the 25^th^ and 75^th^ percentiles, respectively. Extending whiskers represent data points within 1.5 times the interquartile range from lower and upper quartile. Red horizontal lines denote median values, red crosses mark outliers. Following the approach as described in **A-C,** relationship analysis between circadian clock/growth parameters and two additional drug sensitivity was analyzed. Corresponding results for GR50 values are displayed in **(D-F)** and for Hill coefficients of the dose-response curve in **(G-1)** Sample size for **A, D, G:** *n=5* cell lines per drug, except for cisplatin, where n=15 cell lines.

**Figure S6.**
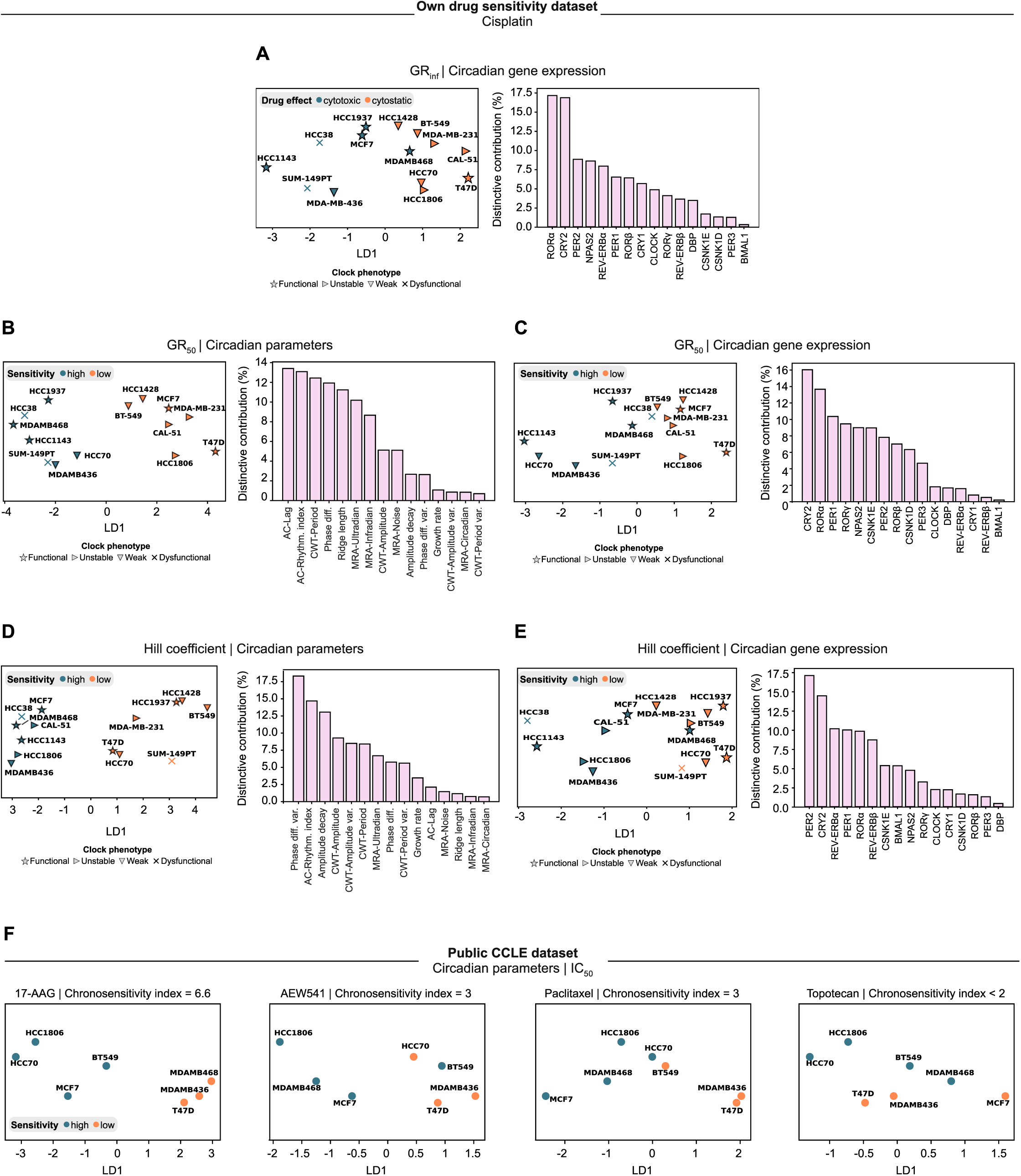
Discriminating between high and low drug sensitivity from circadian parameters and gene expression levels. **(A-E)** Linear discriminant analysis (LDA) on median-binarized drug sensitivity values to cisplatin, as indicated in the plots, using either circadian gene expression data **(A, C** and **E,** n=16 core clock genes), or circadian oscillation parameters **(B** and **D,** n=15 parameters, averaged *Bmal1-Per2* data) as predictors. Cell models with drug sensitivity values below or above the median are colored in green and orange, respectively. The right bar plots rank the individual contribution of each input parameter to the obtained discriminative information. (F) LDA profiles of different drugs, exemplifying distributions for varying chronosensitivity indices, sorted from highest (left panel) to lowest (right panel).

